# Ecological Filtering is a Better Predictor of Microbiome Assembly than Coevolution in Marine Sponges

**DOI:** 10.64898/2026.02.08.704580

**Authors:** Joëlle van der Sprong, Vani Tirumalasetty, Simone Schätzle, Oliver Voigt, Dirk Erpenbeck, Gert Wörheide, Sergio Vargas

**Affiliations:** Department of Earth and Environmental Sciences, Palaeontology and Geobiology, Ludwig-Maximilians-Universität München, Munich, Germany; GeoBio-Center, Ludwig-Maximilians-Universität München, Munich, Germany; SNSB–Bavarian State Collections of Palaeontology and Geology, Munich, Germany

**Keywords:** Sponge Microbiome, Phylogenomics, Phylosymbiosis, Haplosclerida, Target Capture Enrichment, Convergence, Host-Specificity

## Abstract

Understanding how microbiomes evolve in concert with their hosts is crucial for understanding the evolutionary dynamics of complex holobionts. Phylosymbiosis describes a pattern in which microbial community relationships parallel host phylogeny. Although often interpreted as evidence for host–microbiome co-diversification, phylosymbiosis can also arise from ecological filtering, a process in which hosts with similar physiology or morphological characteristics select for similar microbial communities independent of host relatedness. Here, we test co-diversification and ecological filtering as alternative explanations for phylosymbiosis in marine sponges (Order Haplosclerida) from the Red Sea. We inferred a robust host phylogeny using 1,153 target-captured genome-wide loci from 144 specimens and characterised their prokaryotic microbiomes via V4 16S rRNA gene amplicon sequencing. Host clade identity explained ∼50% of the microbiome. However, topological analyses revealed high incongruence between host phylogeny and microbial community composition, and phylogenetic distance was only weakly correlated with microbiome dissimilarity. Partial Mantel tests confirmed that geography did not affect this signal. Trait-based analyses comparing high versus low microbial abundance (HMA–LMA) phenotypes, that are linked to sponge filtration physiology, revealed that this classification was a better predictor of microbiome composition than phylogenetic distance. Furthermore, phylogenetically distant clades with convergent HMA characteristics harboured similar microbiome profiles but had unique amplicon sequence variant (ASV) compositions, with between-trait dissimilarity exceeding within-trait dissimilarity. This demonstrates that sponge–microbiome associations are more influenced by ecological filtering than by host–microbe co-diversification, and suggests that sponge holobionts may be better understood as ecological consortia than as coevolved units.

## Introduction

Phylosymbiosis describes a pattern in which microbial community composition parallels host phylogeny [1]. This concept has gained attention because microbial communities often play critical functional roles in their hosts. For example, gut microbiota influence immunity in mammals [2]; symbiotic bacteria modulate behaviour in social insects [3]; and root-associated communities contribute to pathogen resistance in plants [4]. These functional dependencies raise questions about the extent to which microbial communities are passively acquired or actively selected and functionally integrated by the host. Phylosymbiosis is often interpreted as evidence for host–microbiome co-diversification [5, 6]. However, it can also arise from ecological filtering or niche differentiation when phylogenetically conserved host traits shape microbial community assembly [1, 7]. Theoretical simulations predict that ecological filtering produces weak but significant phylosymbiotic signals, characterised by low host–microbiome correlations and high topological incongruence between host and microbiome dendrograms [7]. However, these predictions have rarely been tested empirically. Distinguishing between these mechanisms is crucial for understanding the evolutionary significance of host–microbiome associations.

Among the various animal groups studied in the context of phylosymbiosis, sponges (Phylum Porifera) stand out for the complexity and stability of their microbial consortia [8, 9]. Sponges are sessile, filter-feeding invertebrates that maintain close associations with diverse microbial communities sometimes constituting up to ∼70% of their dry biomass [9–12]. Within the sponge holobiont, these associations facilitate nutrient cycling, the production of bioactive secondary metabolites, support chemical defence mechanisms [13], and sometimes even biomineralisation [14–16]. Sponges are broadly classified into two categories based on their microbial load and composition. Sponges with high microbial abundance (HMA) phenotypes harbour dense, often sponge-specific microbial communities within their tissues, whereas low microbial abundance (LMA) sponges maintain microbial densities comparable to the surrounding seawater [13, 17]. This HMA–LMA dichotomy is associated with distinct physiological and ecological strategies, including differences in pumping rates, nutrient acquisition, and secondary metabolite production [18, 19]. Furthermore, the HMA phenotype has evolved convergently in multiple sponge lineages [5], complicating interpretations of whether microbiome similarity among HMA sponges stems from shared ancestry or convergent trait-based filtering.

As one of the earliest-branching metazoan lineages [20, 21], sponges have maintained stable microbial associations throughout evolutionary time [22], and a better understanding of how phylosymbiosis structures these partnerships can provide insights into the origins of early animal–microbe coevolution. Thus far, the functional roles of sponge-associated microbes have been well recognised and studied [8, 19, 23]. However, the evolutionary processes shaping these sponge–microbe relationships remain to be further explored. Sponges pump large volumes of seawater (up to thousands of times their body volume per day), providing ample opportunity for horizontal symbiont acquisition [24]. Nevertheless, sponges are not indiscriminate in their associations and can selectively retain beneficial microbes while rejecting foreign or potentially harmful strains [24, 25]. While evidence for phylosymbiosis in sponges exists [24, 26, 27], it is based chiefly on comparisons of microbiomes between distantly related species or supra-specific taxa, often even belonging to different orders or classes [9, 26, 28]. This is problematic given that sponge taxonomy has historically relied on morphological characters that are often unsuitable to unravel true evolutionary relationships [29, 30], leading to the conflation of phylogenetic signal and convergent characteristics.

In this study, we focus on the species-rich and systematically complex demosponge order Haplosclerida. Demospongiae represents the largest and most diverse sponge class, comprising approximately 80% of all described sponge species and encompassing the majority of studied sponge–microbe associations [31]. Recent genome-scale phylogenetic analyses (i.e., phylogenomics) have resolved the evolutionary relationships within Haplosclerida [30, 32], but the current taxonomy remains based mainly on morphological characters that conflict with molecular hypotheses [33–35]. Using target-capture enrichment of clade-specific elements (CSEs), exon loci optimised for resolving phylogenetic relationships within Haplosclerida [30, 32], we inferred a robust phylogeny of haplosclerid specimens collected along the Arabian coastline of the Red Sea. This geologically isolated basin is characterised by distinctive latitudinal gradients in temperature, salinity and nutrient availability, and represents a marine biodiversity hotspot known for high levels of endemism [36]. If phylosymbiosis is explained by coevolutionary processes, microbial community composition is expected to closely mirror host phylogenetic relationships, regardless of phenotypic traits that affect ecological filtering. Alternatively, if ecological filtering drives these patterns, functional traits such as HMA–LMA phenotypes should better predict microbiome composition than phylogenetic distance. To test these competing hypotheses, we combined a genome-scale host phylogeny with 16S rRNA amplicon sequencing of sponge prokaryotic communities spanning the phylogenetic breadth of the Haplosclerida. Combining a phylogenetically robust inference with replicated sampling of the same species across a geographically constrained region enabled us to determine the relative contributions of host phylogeny, functional traits, and environmental variation to structuring microbiome assembly. Here, we reveal that HMA–LMA status is a better predictor of microbial dissimilarity than host phylogeny, and that geography explained little of the observed variation. Consequently, sponge holobionts may be better conceptualised as ecological consortia assembled by functional compatibility than as tightly coevolved units.

## Materials and Methods

### Sample collection

Haplosclerid specimens (n = 162) were collected along the Arabian coastline of the Red Sea between 2012 and 2017 as part of the Red Sea Biodiversity Project (Senckenberg Research Institute; King Abdulaziz University) and sampling efforts by King Abdullah University of Science and Technology (KAUST) [32, 37]. Specimens were collected by hand-picking in intertidal zones, SCUBA diving, or trawling at depths of 1–63 m. Samples were sorted, photographed, and preserved in 95% ethanol.

### DNA extraction, library preparation, and target capture enrichment

Samples were processed according to the protocol described by Van der Sprong et al. (2024). DNA was extracted using the NucleoSpin® Tissue Kit (Macherey-Nagel) and quantified using a Qubit 2.0 fluorometer. Libraries were prepared using the xGenTM ssDNA and Low-Input DNA Prep kit at half the manufacturer’s recommended reaction volumes. Libraries were quantified and quality-controlled using a Bioanalyzer 2100, and amplified using KAPA HiFi polymerase to provide 250 ng for hybridisation. Target capture was performed using a custom multilocus probe set consisting of 20,000 baits targeting 2,956 exon regions (i.e., clade-specific elements) across Haplosclerida [30, 32] with pools of eight libraries following the MyBaits target enrichment protocol. Target-enriched libraries were sequenced in paired-end mode (2 × 150 bp) on an Illumina MiniSeq platform in high-throughput mode.

### Host phylogenomic analysis

Demultiplexed Illumina raw reads were quality-controlled and trimmed using *fastp* [38]. Cleaned reads were processed using a modified PHYLUCE v1.7.3 workflow [32, 39, 40]. Reads were assembled with SPAdes, and CSE loci were selected based on a minimum 85% identity and coverage between assembled contigs and exon capture probes. Selected CSE loci were extracted and aligned, followed by internal trimming with GBlocks using default parameters (as implemented in PHYLUCE). The final CSE loci alignment was concatenated and formatted to PHYLIP format. Maximum likelihood (ML) phylogenetic inference was performed using RAxML v8.2.12 [41] with rapid bootstrapping (GTR+GAMMAX model, 1000 bootstrap replicates).

### 16S rRNA amplification and sequencing

The V4 hypervariable region of the 16S rRNA gene was amplified using Illumina-ready primers 515F and 806R [42]. PCR reactions (25 µL total volume) contained 12.5 µL GoTaq® G2 Hot Start Master Mix 2× (Promega), 1 µL each primer (5 µM), 1 µL template DNA (5 ng µL⁻¹), and 9.5 µL nuclease-free water. Amplification conditions were 95°C for 3 min; 35 cycles of 95°C for 30 s, 55°C for 30 s, 72°C for 30 s; 72°C for 5 min. Amplicon quality was verified by agarose gel electrophoresis (1.5%, 120 V, 30 min). Bands (∼380 bp) were excised and purified using the NucleoSpin Gel and PCR Clean-up kit (Macherey-Nagel), eluted in 20 µL nuclease-free water, and quantified using a Qubit 2.0 fluorometer. Amplicons were sequenced in paired-end mode (2 × 150 bp) on an Illumina MiniSeq platform using the MiniSeq Mid Output Reagent Kit (300 cycles) with dual-indexed primers, as described by Pichler et al. (2018).

### 16S sequence processing and quality control

Raw sequencing data were processed using VSEARCH v2.5.0 [43]. Paired-end reads were merged (minimum overlap 45 bp, zero mismatches), quality filtered (maximum expected error ≤ 0.5, length 225–275 bp, no ambiguous bases), and dereplicated per sample with singletons removed. Chimeric sequences were removed using UCHIME with both de novo and reference-based methods against the UCHIME gold database [44].

Clean reads were further processed using DADA2 v1.8 [45] in R v4.5.2. Of 162 initial samples, 14 were excluded due to low sequencing depth (< 5,000 reads) and 4 due to missing host phylogenetic data, leaving 144 for downstream analysis. Reads were quality filtered with a maximum expected error of two for both forward and reverse reads, quality score truncation at 2, no ambiguous bases allowed, and PhiX contamination removed. Error rates were estimated from filtered data, and amplicon sequence variants (ASVs) were inferred independently for forward and reverse reads using the DADA2 denoising algorithm before merging paired reads. Chimeric sequences were identified and removed using the consensus method in *removeBimeraDenovo*. Taxonomic classification was performed against SILVA v138.1 [46] using both the nr99 training set and species assignment database for species-level assignments where possible. ASVs that could not be classified at the Kingdom or Phylum level were validated using BLASTn against the NCBI nucleotide database. Sequences assigned to mitochondrial or chloroplast lineages, or lacking significant prokaryotic hits, were removed.

### Haplosclerid holo-microbiome composition analysis

Phylosymbiosis analyses were performed in R v4.5.2. ASV count tables, taxonomy assignments, and phylogenetic trees were imported into phyloseq objects [47]. Samples with < 5,000 reads (n = 7) were excluded. The filtered dataset was transformed to compositional abundances (microbiome package) for beta-diversity analyses [48]. Bray–Curtis dissimilarity matrices were calculated using *phyloseq*. Non-metric multidimensional scaling (NMDS; 100 iterations) and principal coordinates analysis (PCoA) were visualized using ggplot2.

Betadispersion (betadisper, vegan) assessed homogeneity of multivariate dispersion among clades, with significance tested using *permutest* (999 permutations) [49]. PERMANOVA (adonis2, vegan; 999 permutations) tested the effects of host clade on microbial composition. Pairwise comparisons were performed using *pairwise.adonis* with the Benjamini–Hochberg correction [50]. Host phylogenetic distances were calculated as cophenetic distances (ape) [51]. Microbial community distances were calculated using Bray–Curtis dissimilarity, weighted UniFrac, and unweighted UniFrac. Mantel tests (vegan; Spearman correlation, 9,999 permutations) assessed correlations between host phylogeny and microbiome dissimilarity. Partial Mantel tests controlled for geographic distance and phylogeny (calculated using Haversine distances, geosphere) [52]. Microbial dendrograms were constructed using UPGMA clustering (*hclust*). Normalised Robinson–Foulds (nRF) distances between host and microbial trees were calculated using *cospeciation* (phytools; 999 permutations) [53]. Tanglegrams were constructed using *dendextend* with step2side untangling [54].

Samples were classified as high microbial abundance (HMA; clades G01, G06, G12) or low microbial abundance (LMA; clades G03–G05, G07, G09, G15, G16) based on previously published characterisations of these taxa [55]. These classifications were consistent with the phylum-level profiles observed in our data, with HMA clades enriched in Chloroflexi and Acidobacteriota, and LMA clades dominated by Cyanobacteria (Figures 1 and 2). Binary trait distance matrices (0 = same type, 1 = different type) were constructed.

**Figure 1.**
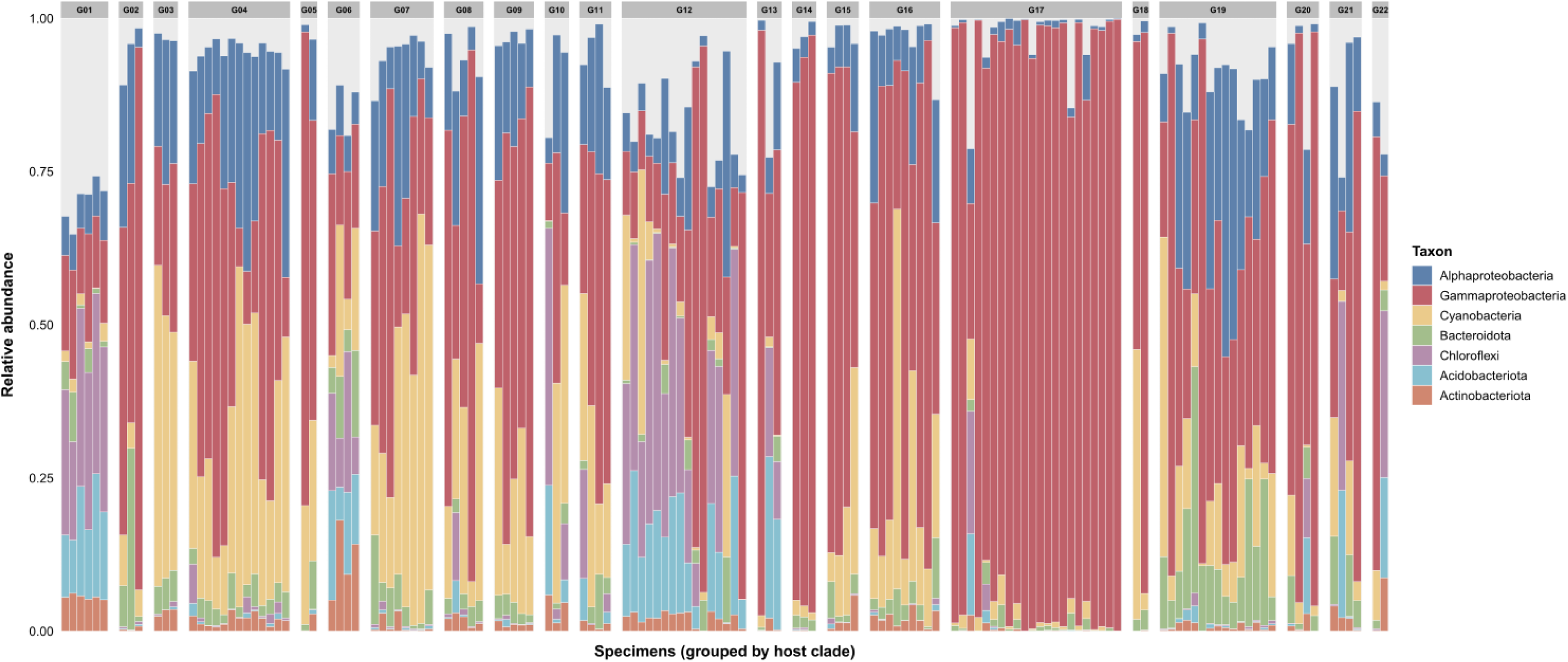
Microbial profiles of Red Sea Haplosclerida sponges. At phylum level, with Proteobacteria subdivided into the classes Alpha- and Gammaproteobacteria. Relative abundances (%) are shown for taxa with > 0.25% mean abundance. The operational host clades (G01-G22) indicated follow phylogenetic order.

**Figure 2.**
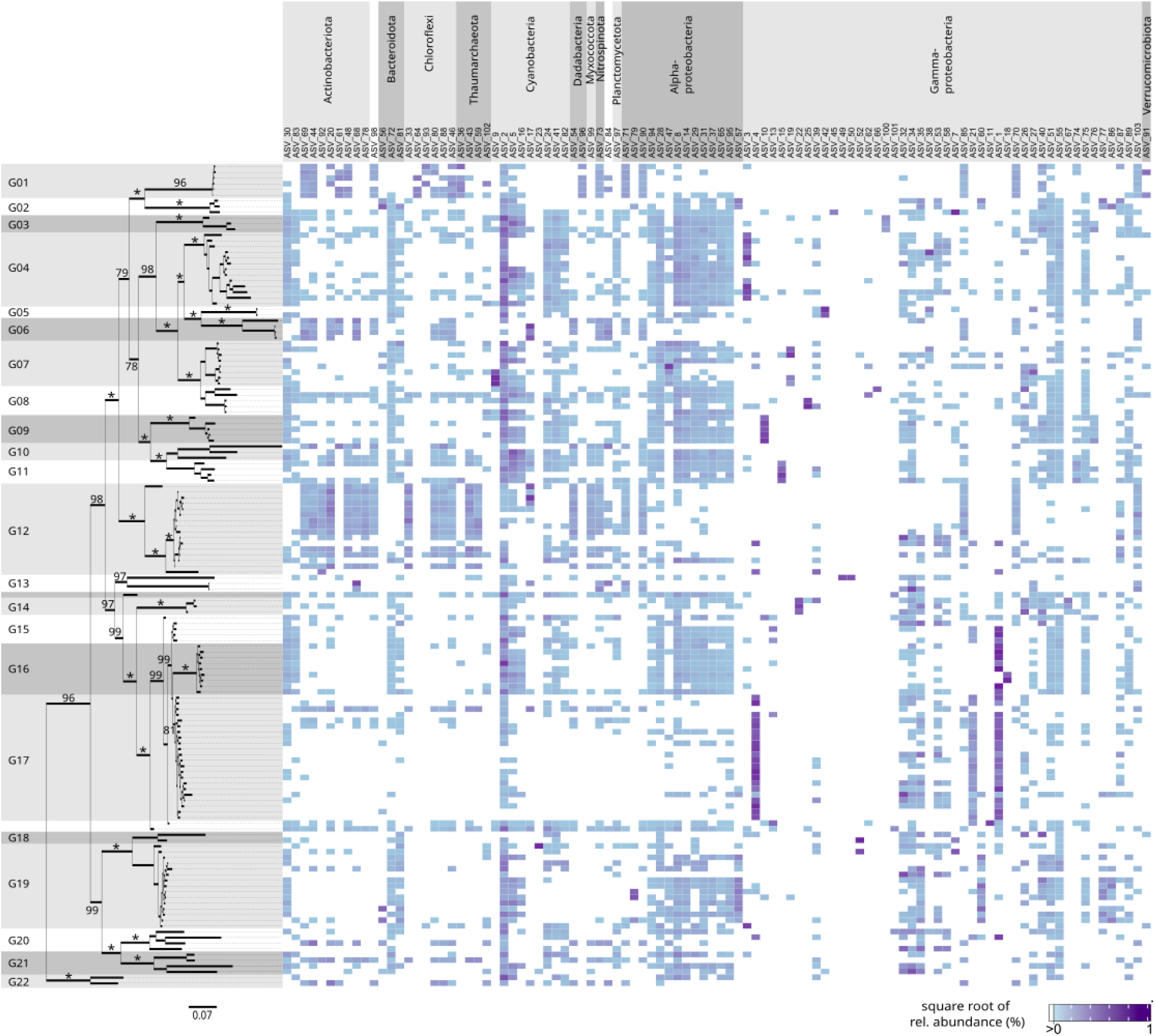
Host phylogeny and amplicon sequence variant (ASV) distribution in Red Sea Haplosclerida sponges. Maximum Likelihood (ML) phylogeny of 144 specimens was based on 1,153 Clade-Specific Element (CSE) loci with ≥ 25% taxon coverage per locus (276,047 bp total alignment). Bootstrap probability (BP) values are shown at nodes; asterisks (*) indicate BP = 100. Scale bar represents substitutions per site. The heatmap shows the mean relative abundances (square-root transformed) of the 100 most abundant ASVs across host clades (G01-G22). ASVs are grouped by phylum, with Proteobacteria subdivided into Alpha-and Gammaproteobacteria. The two ASVs ASV_98 (AncK6) and ASV_84 (PAUC34f; uncultured sponge bacterium) could not be confidently classified at phylum level.

Mantel tests compared trait similarity versus phylogenetic distance as predictors of microbiome dissimilarity. PERMANOVA quantified trait effects on HMA–LMA subsets. Variance partitioning used sequential PERMANOVA with PCoA-transformed phylogenetic and geographic distance matrices (3 axes each) to decompose microbiome variance into trait, phylogeny, geography, and shared components. Pairwise Bray–Curtis dissimilarities were classified as within-HMA, within-LMA, or between HMA-LMA comparisons. Kruskal–Wallis tests (*kruskal.test*) assessed group differences, with post-hoc pairwise comparisons using Dunn tests (FSA) with Bonferroni correction [56]. All analysis scripts are available at: https://github.com/PalMuc/RedSeaHaploscleridaPhylosymbiosis.

## Results

### Host phylogenomics

Of 162 haplosclerid specimens collected from the Red Sea, 144 passed quality control for downstream analyses: 14 were excluded for insufficient sequencing depth, 4 for missing host data (Supplementary Table 1). Target capture CSE loci retrieved a mean of 2,147,166 ± 1,698,472 SD clean reads per sample (range: 220,342–10,926,406 reads). The number of captured CSE loci per specimen averaged 628 ± 246 SD (range: 127–1,158 loci).

ML phylogenetic inference of the 144 haplosclerids was based on 1,153 CSE loci with ≥ 25% taxon coverage per locus (Supplementary Figure 1). Based on the inferred phylogeny, we selected 22 operationally distinct genetic clusters (G01–G22). Bootstrap support values were high (BP ≥ 95%) for most nodes. Of the 144 specimens analysed, 23 had previously been identified to species or (sub)genus level based on morphological characteristics [32]. However, morphological identification in Haplosclerida is problematic due to a paucity of diagnostic traits, phenotypic plasticity, widespread convergence, and often contradicts molecular phylogenetic hypotheses, with almost all supraspecific assignments known to be non-monophyletic [30, 33, 35]. Therefore, specimens in this study were primarily analysed based on their molecular phylogenetic relationships.

### Microbial assemblages

Sequencing of the V4 region of the 16S rRNA gene amplicon resulted in an average of 23,321.67 ± 22,404.09 SD clean reads per sample (range: 4,981–212,332). The final dataset comprised 9,018 amplicon sequence variants (ASVs) across 144 sponge specimens (Supplementary Table S1). Following ASV inference, chimaera removal, and exclusion of mitochondrial and chloroplast sequences, seven additional sponge specimens fell below the 5,000-read threshold. Rarefaction analysis indicated that microbial diversity was undersampled in these samples. The final dataset comprised 8,807 ASVs across 136 specimens, with read depth ranging from 5,173 to 143,699 reads (median: 14,629; mean: 20,157).

### Clade-specific microbiome composition and diversity

Microbial community composition varied among the 22 operational host clades (Figure 1), and closely related host clades did not necessarily harbour similar microbial community profiles. Alpha- and Gammaproteobacteria were the most abundant groups across all clades, though their relative abundances differed. Several clades were characterised by high abundances of Gammaproteobacteria (e.g., G14, G15 and G17), while others were more or equally dominated by Cyanobacteria (e.g., G03–G05, G07–G11 and G12). Clades G01, G06 and G12 were characterised by high relative abundances of the phyla Chloroflexi and Acidobacteriota. The phyla Bacteroidota, Chloroflexi, Acidobacteriota and Actinobacteriota were present in most clades, but generally at lower abundances.

The distribution of the 100 most abundant ASVs against the host phylogeny demonstrated host-specificity of the microbial assembly (Figure 2), and their patterns appeared much more clade-specific. Gammaproteobacteria ASVs (e.g., ASV_4: Burkholderiales, EC94) were found to be highly abundant in clade G17, as was Nitrosococcaceae ASV_1 (AqS1), which was also highly abundant in clade G16. In other host clades, these ASVs were much less abundant or absent. Similarly, ASVs belonging to the phyla Actinobacteriota, Chloroflexi and Thaumarchaeota were found in clade G12, with partial overlap observed in clades G01 and G06. The ASV representing the Acidobacteriota phylum (ASV_92: Thermoanaerobaculaceae, Subgroup 10) was abundant in clade G12, but rare or absent in almost all other clades. Furthermore, consistent with the observations at the phylum level (Figure 1), the G03–G05, G07–G11 and G14–G16 clades harboured high abundances of multiple Cyanobacteria ASVs. Several ASVs were detected across most or all host clades, suggesting they are potential core members of the microbiome.

### Phylosymbiosis analysis

NMDS analysis of Bray–Curtis dissimilarities produced a stress value of 0.21, indicating limited two-dimensional representation (Supplementary Figure 2). PCoA revealed better community structuring (Figure 3), with several clades forming discrete clusters (G12, G15, G16, G17) while the other clades showed considerable overlap. PERMANOVA indicated that host clade explained approximately half of the variation in microbial community composition (R^2^ = 0.496, p = 0.001). Betadispersion analysis showed marginally non-significant differences in within-clade dispersion (F = 1.72, p = 0.052), confirming that the PERMANOVA results cannot be the result of unequal dispersion between the clades.

**Figure 3.**
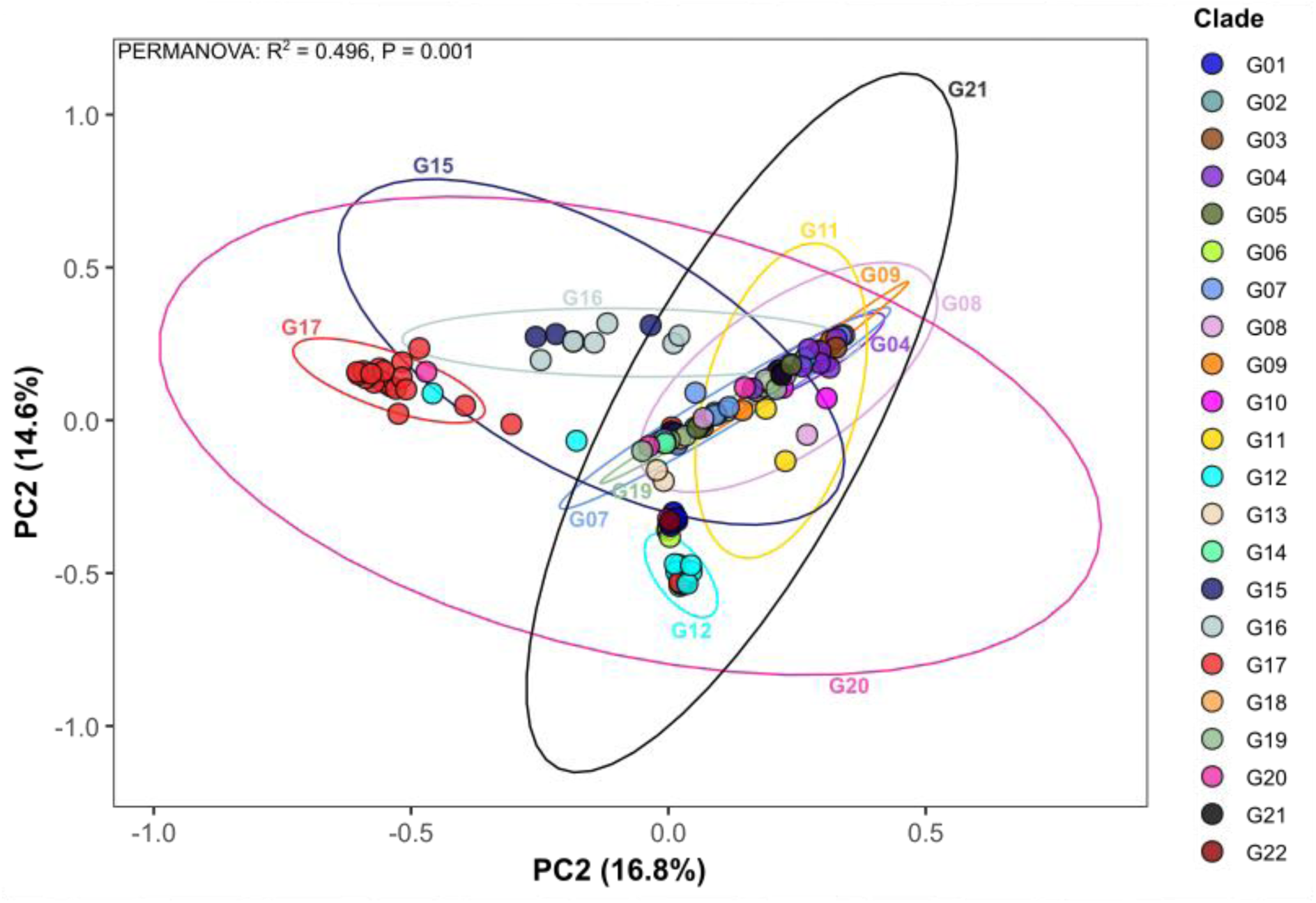
Principal coordinates analysis (PCoA) of microbial community composition in Red Sea Haplosclerida. Ordination based on Bray-Curtis dissimilarities of compositionally transformed transformed ASV abundances. Each point indicates an individual specimen, coloured by host clade assignment (G01–G22). Ellipses indicate 95% confidence intervals.

Mantel tests revealed significant but weak correlations between host phylogenetic distances and microbial dissimilarities across all beta-diversity metrics (Table 1A). Normalised Robinson–Foulds (nRF) distances exceeded 0.89 for all metrics, indicating some degree of topological incongruence between host phylogeny and microbiome dendrograms, even though this was significantly different from random expectations. Entanglement values were low (< 0.28), suggesting partial congruence in tip order between trees (Supplementary Figure 3). The phylogenetic correlation remained largely unchanged after geographic correction, indicating that spatial autocorrelation does not confound a signal of phylosymbiosis. Geographic distance itself showed minimal correlation with microbiome dissimilarity (Table 1B; Supplementary Figure 4).

**Table 1.**
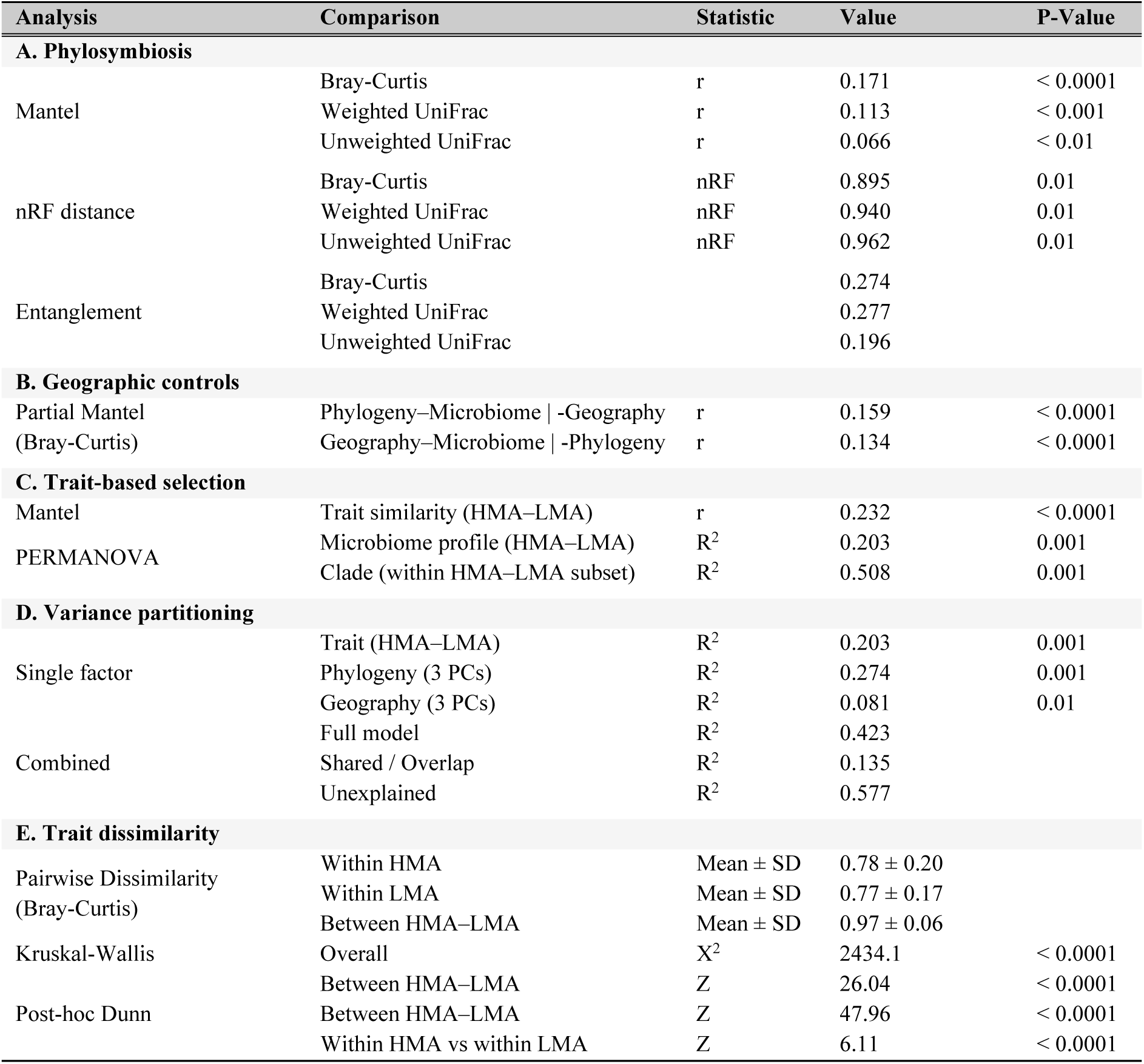
Statistical tests of phylosymbiosis and trait-based selection in Red Sea Haplosclerida. Mantel correlation coefficients (r) between host phylogenetic distances and microbial dissimilarities (Spearman, 9,999 permutations). Partial Mantel tests control for geographic or phylogenetic distance. Normalised Robinson–Foulds (nRF) distances between host tree and microbial dendrograms (999 permutations; 0 = identical, 1 = different). Entanglement values measure tip order congruence (0 = good fit, 1 = misalignment). PERMANOVA R² values indicate proportion of variance explained (999 permutations). Variance partitioning based on sequential PERMANOVA with PCoA-transformed distance matrices (3 PCs). Pairwise dissimilarities (Bray-Curtis) were classified by trait comparison type; differences assessed using Kruskal-Wallis test with post-hoc Dunn tests (Bonferroni-corrected p-values). High microbial abundance (HMA) clades; Low microbial abundance (LMA) clades.

### Trait-based microbiome filtration

Trait similarity (HMA–LMA) was a better predictor of microbiome dissimilarity than phylogenetic distance (Mantel r = 0.232 vs r = 0.171; Table 1C). HMA–LMA signatures explained 20.2% of microbiome variation; however, clade identity still held significant explanatory power (50.8%), even within HMA–LMA subsets. Variance partitioning revealed that trait, phylogeny, and geography collectively explained 42.3% of microbiome variation (Table 1D). A large proportion of the variance (13.5%) was shared, consistent with the phylogenetic conservation of HMA–LMA traits within lineages. However, most of the variance (57.7%) remained unexplained. Between-trait dissimilarities (HMA vs LMA) were significantly higher than within-trait comparisons, with all pairwise differences highly significant (Table 1E). Within-HMA and within-LMA dissimilarities were comparable (0.78 vs 0.77), indicating similar levels of compositional variation within each functional category.

## Discussion

### Phylosymbiosis does not indicate coevolution

Lim and Bordenstein (2020) defined phylosymbiosis as a pattern in which the relationships among microbial communities recapitulate host phylogeny. Following this definition, symbiosis is interpreted in the broadest sense. It includes any kind of association between organisms of different species, independent of the duration or functional nature of the interaction. Therefore, phylosymbiosis is mechanism agnostic only indicating only the presence of a pattern or correlation. It is not, in itself, a direct result of co-diversification, vertical transmission, or mutualistic interactions. It can originate through various processes, such as cospeciation, the horizontal transmission of phylogenetically conserved host traits, or the ecological filtering of environmental microbes each generation [1].

Our results confirm the presence of phylosymbiosis in Red Sea Haplosclerida, but make a critical distinction between host-specific microbiome structuring and coevolutionary microbiome assembly. PERMANOVA analysis showed that host clade identity accounted for nearly half of the variation in microbial community composition, indicating host-mediated influence on microbiome composition. However, the extent to which this host effect is observed is not proportional to phylogenetic relatedness. Mantel tests revealed only weak correlations between host phylogenetic distance and microbiome dissimilarity across all beta-diversity metrics (Table 1A), and topological comparisons showed high incongruence between host phylogeny and microbial dendrograms (Supplementary Figure 3). Partial Mantel tests confirmed that this weak phylogenetic signal was unaffected by geography, as the correlation changed only slightly following spatial correction (Table 1A–B). Thus, even though host identity is an important factor in microbiome composition, phylogenetic distance between hosts is a poor predictor of microbiome dissimilarity.

### Ecological filtering as a more parsimonious explanation

The combination of host-microbiome specificity and weak phylogenetic signal is a consistent finding across the literature on sponge microbiomes [5, 9, 24, 27]. For example, Thomas et al. (2016) reported that host phylogeny explained only 5% of additional variance after accounting for host identity, and similar patterns of high topological incongruence despite apparent species-specific effects have been observed in Petrosiidae and Caribbean sponges [5, 27]. These results are often presented or interpreted as evidence for coevolutionary host-microbiome relationships [5, 6, 57]. However, beta-diversity clustering alone is insufficient to confirm that phylogeny is the primary driver of microbiome composition without explicit tests for correlations between host phylogenetic distance and microbiome dissimilarity [7]. Variables such as geography, depth, or incomplete taxon sampling are usually invoked to explain weak effect sizes, preserving the assumption that coevolution underlies these patterns.

Our study challenges this interpretation because phylosymbiosis can arise from ecological filtering, independent of co-diversification. We find that HMA–LMA status acts as the key factor shaping the microbiome, with hosts of similar filtration physiology harbouring similar microbiomes irrespective of phylogenetic relatedness. Phylosymbiotic patterns may arise as a by-product of ecological filtering, because host traits that influence microbiome composition often are shared within lineages. This mechanism has been demonstrated theoretically [7], but sponge microbiome studies have rarely explicitly tested their findings against filtering predictions, such as weak Mantel correlations and high topological incongruence. This distinction is not limited to sponges, as phylosymbiotic patterns across the tree of life are often interpreted as evidence for co-diversification by default without the mechanistic evidence to support this. Under a filtering scenario, simulations predict weak Mantel correlations (r ≈ 0.2) and high topological incongruence (nRF ≈ 0.9) [7]. Our empirical results fall within these predictions (Table 1), as do those from other sponge microbiome studies (e.g., Van der Windt et al. 2025: r = 0.202; nRF > 0.84). If coevolutionary processes were the main driver behind the phylosymbiosis pattern, we would expect higher Mantel correlations and greater topological congruence between host phylogeny and microbiome composition. By focusing on a single taxonomic group within a single biogeographic region, we were able to control for many conflicting factors, such as deep phylogenetic divergence, large-scale environmental gradients, and geographic distance. Nevertheless, the phylogenetic signal remained weak. Therefore, a more parsimonious explanation is that sponge microbiome assembly is driven by host-specific ecological filtering rather than evolutionary conservation.

Hosts possess traits that influence the acquisition and retention of microbes, such as an aquiferous system architecture, biochemical environment, or immune-like recognition systems, which vary among host species and shape prokaryotic community composition [8, 19]. When specific traits are phylogenetically conserved within host lineages, statistical correlations with microbiome composition are likely to emerge as a result of phylogenetic inertia, producing a phylosymbiotic signal without necessarily requiring co-diversification. This is also supported by earlier studies demonstrating that microbiome dissimilarity in sponges is mainly driven by differential abundances of shared taxa, rather than the presence of unique, host-specific microbial consortia [9, 26]. Easson and Thacker (2014) showed that half of the sponges studied had no species-specific microbial taxa, despite near-perfect host clustering (R² = 0.735). If coevolutionary processes were mainly responsible for this, the core microbiome would be dominated by more host-specific specialists.

### Convergent evolution of HMA–LMA phenotypes

Our analysis reveals that convergent host phenotypes produce similar microbiome compositions despite distant phylogenetic relationships (Figures 1 and 2). Clades G01 (*Chalinula*), G06 (*Neopetrosia*), and G12 (*Xestospongia*) are phylogenetically distant. However, they each evolved HMA phenotypes independently within the Haplosclerida [55]. These clades share similar prokaryotic enrichment patterns, with relatively high abundances of Chloroflexi and Acidobacteriota, as expected for HMA sponges [17]. HMA symbiosis has evolved at least 21 times independently from ancestral LMA states across sponges, with nearly half of these transitions (9/21) occurring within Haplosclerida alone [5]. Furthermore, frequent reversals from HMA to LMA states (7/8) indicate evolutionary lability instead of stable co-evolved partnerships. Although Pankey et al. (2022) acknowledged that the convergent nature of HMA communities could support a role for environmental filtering, they nevertheless interpreted their findings as evidence for coevolution (i.e., co-diversification between host and microbial lineages, inferred from cophylogenetic patterns). This interpretation is difficult to reconcile not only with our findings, but also with the broader patterns documented across studies [9, 26, 27, 58], including Pankey et al.’s own observation of multiple independent HMA origins and frequent HMA-to-LMA reversals. Given that HMA microbiomes are assumed to represent coevolved partnerships, frequent reversals are expected to be rare, and hosts transitioning to LMA morphology would be expected to lose their co-diversified microbial partners. Moreover, co-diversification predicts that each independent HMA lineage will develop unique microbial partnerships through separate evolutionary trajectories and be less likely to recruit similar assemblages consistently.

### Mechanisms of ecological filtering

The convergence of HMA phenotypes described above provides the context for understanding how ecological filtering operates at the mechanistic level. HMA phenotypes possess dense mesohyl, reduced aquiferous systems, and slower pumping rates [17, 18, 59], resulting in low water turnover that likely favours slow-growing bacterial lineages over fast-growing opportunists [60]. Furthermore, such dense tissue organisation has been observed to create steep oxygen gradients [61, 62], which benefit prokaryotes that thrive under low-oxygen or anaerobic conditions. The HMA clades in our dataset (e.g., G01, G06, G12) share these characteristics and have comparable enrichment patterns of Chloroflexi and Acidobacteriota despite their phylogenetic distance. LMA phenotypes, on the other hand, possess thinner tissue, higher pumping rates, and less dense microbial communities [59], with higher relative abundances of Alpha- and Gammaproteobacteria [17, 63]. Their branching, encrusting, or thin-walled body structures permit light penetration, supporting photosynthetic symbionts such as cyanobacteria [64]. Clades G03–G05 and G07–G11 in our dataset show these LMA-typical profiles, with cyanobacterial dominance and lower overall microbial diversity (Figures 1 and 2). These HMA–LMA phenotype differences are linked with distinct trophic strategies. HMA sponges derive 71–93% of their carbon from dissolved organic matter (DOM), whereas LMA sponges obtain only 0–5% from DOM and are more reliant on particulate organic matter and bacterioplankton [59]. Hence, HMA sponges are more likely to favour heterotrophic microbial communities capable of processing DOM. LMA sponges, on the other hand, support phototrophic symbionts that supplement host nutrition by fixing carbon.

Experimental evidence demonstrated that HMA sponges resist colonisation by external coral-derived bacteria, while LMA sponges readily incorporated them [65]. This differential uptake likely relates, at least in part, to phenotypic differences between HMA and LMA sponges. However, phenotype alone may not fully explain colonisation resistance. The established microbial community itself may contribute as well, either through direct competition, the production of antimicrobial compounds, or the occupation of available metabolic niches [19, 66]. In addition, sponges appear to function as relatively closed systems compared to other benthic organisms. Sponge-associated microbiomes were found to have the highest dissimilarity to ambient seawater among reef holobionts, suggesting that sponges actively select their microbial partners [67]. Both host phenotype and microbiome-mediated effects are expected to drive host-specific community assembly.

### Functional redundancy and metabolic integration

Ecological filtering explains the host-specificity of their microbial consorts, but not why these communities resist colonisation once established. Two processes may contribute to this stability. First, functional redundancy, whereby multiple taxa perform overlapping metabolic functions, and second, tight metabolic integration among community members. Fan et al. (2012) demonstrated that sponge microbiomes achieve equivalent metabolic outcomes through analogous pathways. For example, different sponge species perform denitrification using distinct enzyme variants (*NarG* versus *NapA*) [28], demonstrating that taxonomically distinct microbiomes can be functionally equivalent. This means that hosts with similar physiological properties select for functionally similar, though not necessarily taxonomically identical, symbionts. Also, horizontal gene transfer can facilitate the spread of host-adaptive functions among microbial community members, enabling functional convergence without shared ancestry. Not all functions are redundantly distributed, however. A distinction can be made between “community functions” spread across diverse taxa (e.g., carbon fixation, B-vitamin synthesis) and “keystone functions” restricted to specific lineages, such as ammonia oxidation by Thaumarchaeota [68]. Engelberts et al. (2020) demonstrated that ammonia oxidation in *Ircinia ramosa* was exclusively performed by Thaumarchaeota despite extensive redundancy for other pathways. This suggests that Thaumarchaeota occupy an obligate niche that alternative taxa cannot replace. The Thaumarchaeota detected in our clade G12 (*Xestospongia*) may also represent keystone taxa for nitrogen cycling. Keystone taxa also show metabolic flexibility under stress. It has been observed that in tolerant sponges Thaumarchaeota upregulate carbon fixation and ammonia oxidation when exposed to ocean acidification, whereas in sensitive sponges this capacity was lost entirely [69].

A third feature of HMA microbiomes is tight metabolic integration among community members. For example, ammonia-oxidising Thaumarchaeota and nitrite-oxidising Nitrospira in *Cymbastela concentrica* were found to be spatially co-localised and metabolically coupled, with host-derived organic nitrogen compounds (creatine, creatinine, urea) fuelling the nitrogen cycle [66]. This functional integration creates a barrier to colonisation as incoming microbes must not only find an unoccupied niche but also integrate into existing metabolic networks. The combination of functional redundancy (stability), keystone taxa (essential functions), and metabolic integration (coordination) may explain why HMA microbiomes maintain distinct compositions across evolutionary time while resisting colonisation by environmental prokaryotes.

### Future directions

Variance partitioning analysis showed that the combined effects of HMA–LMA phenotypes, host phylogeny, and geography explained only 42.2% of microbiome variation, leaving 57.8% unexplained. Several factors may contribute to this and need further investigation. We acknowledge that taxonomic composition alone does not fully capture the complexity of host-microbiome relationships. Different microbial taxa can converge on similar metabolic capacities through functional redundancy [28, 70, 71], and shifts in the relative abundance of shared taxa can alter functional output even when taxonomic profiles appear similar [69].

Integrating metagenomic and metatranscriptomic approaches could reveal whether functional profiles show patterns not apparent from taxonomic composition and whether these profiles are better explained by host phylogeny, host traits, or environmental factors [58]. Also, local environmental filtering conditions at the microhabitat level remain largely uncharacterised.

Factors such as water flow regimes, light penetration, proximity to other benthic organisms, temporal dynamics, and sedimentation exposure influence microbial community assembly but are challenging to assess systematically across large-scale sampling efforts. Depth has been shown to affect sponge microbiome composition in other systems [27, 57, 72], but we were unable to test for this effect due to incomplete depth records for our specimens (Supplementary Table S1), though most of our specimens occurred in relatively shallow waters (< 20 m). Finally, sponges harbour diverse macro-organisms, such as polychaetes and crustaceans [73–75], whose influence on microbiome composition remains largely unexplored. Recent work detected secondary metabolites produced by associated macro-fauna [72], suggesting that macro-symbionts influence the chemical environment within their hosts, but the consequences for microbiome assembly are unknown. Because phylosymbiosis may not be explained solely by coevolution, we encourage future studies to focus on a broader range of mechanisms that could drive these patterns as well. This will provide more insights into the factors shaping microbial community composition in holobionts and how these systems adapt to changing environmental conditions.

## Acknowledgements

We thank Gabriele Büttner and Nora Dotzler for laboratory assistance. Sample collection was facilitated through the Red Sea Biodiversity Project, a collaboration between King Abdulaziz University, King Abdullah University of Science and Technology, and the Senckenberg Research Institute (Grant No. I/1/432-DSR).

## Data Accessibility and Benefit-Sharing

Raw sequence data are deposited in GenBank under Bioproject accession no. PRJEB102953. All analysis scripts are available at: https://github.com/PalMuc/RedSeaHaploscleridaPhylosymbiosis and https://doi.org/10.5281/zenodo.18523495.

## Author Contributions

Conceptualisation: SV, DE, GW. Investigation: VT, JS, SS. Formal analysis: JS, SV. Resources: DE, GW, OV. Writing of the original draft: JS. Writing of the review and editing: SV. All authors reviewed the manuscript and agreed with its contents.

