## Supplementary figures and images for "Ecological Filtering is a Better Predictor of Microbiome Assembly than Coevolution in Marine Sponges"

### Figure S1

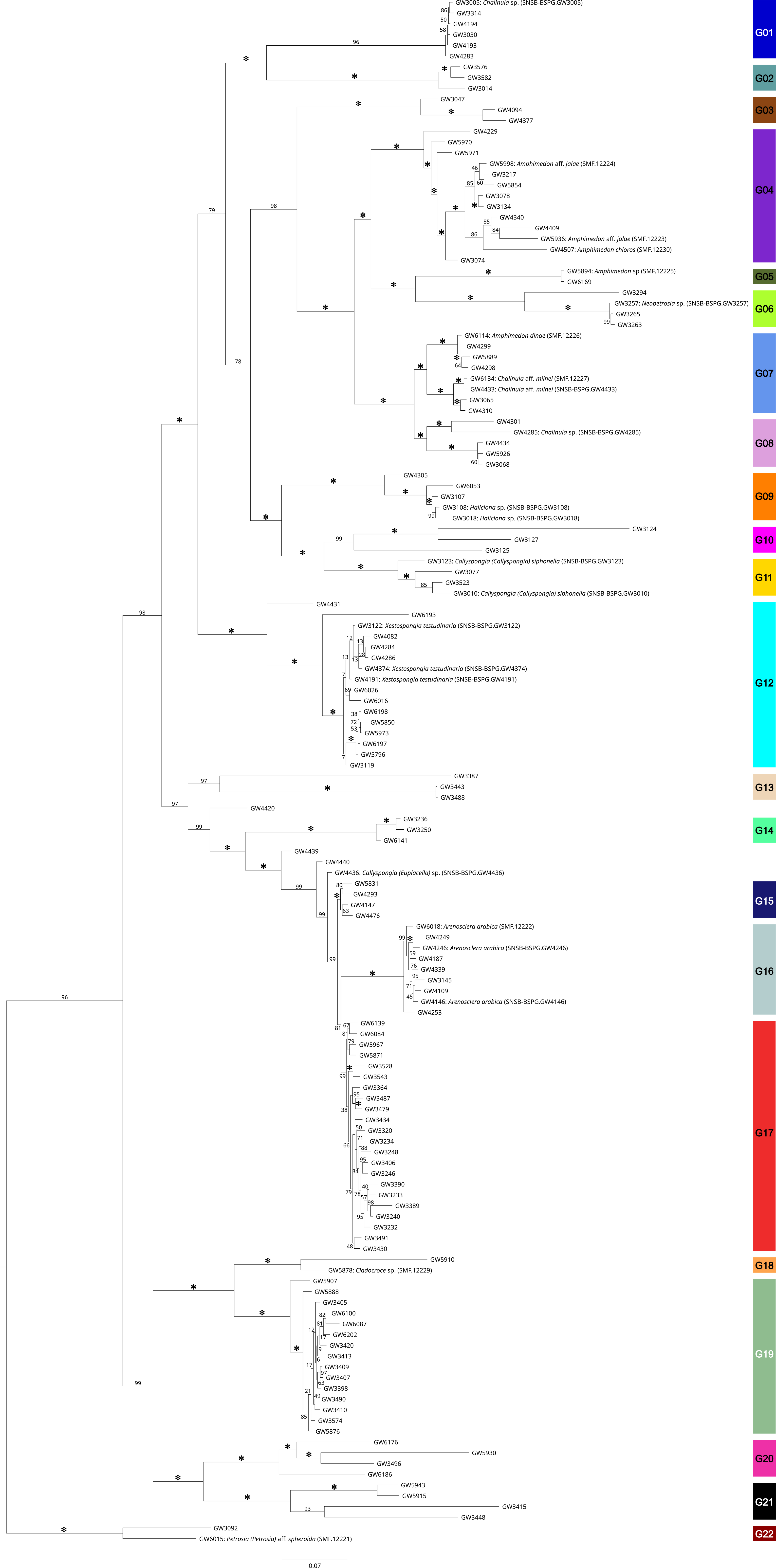

### Figure S2

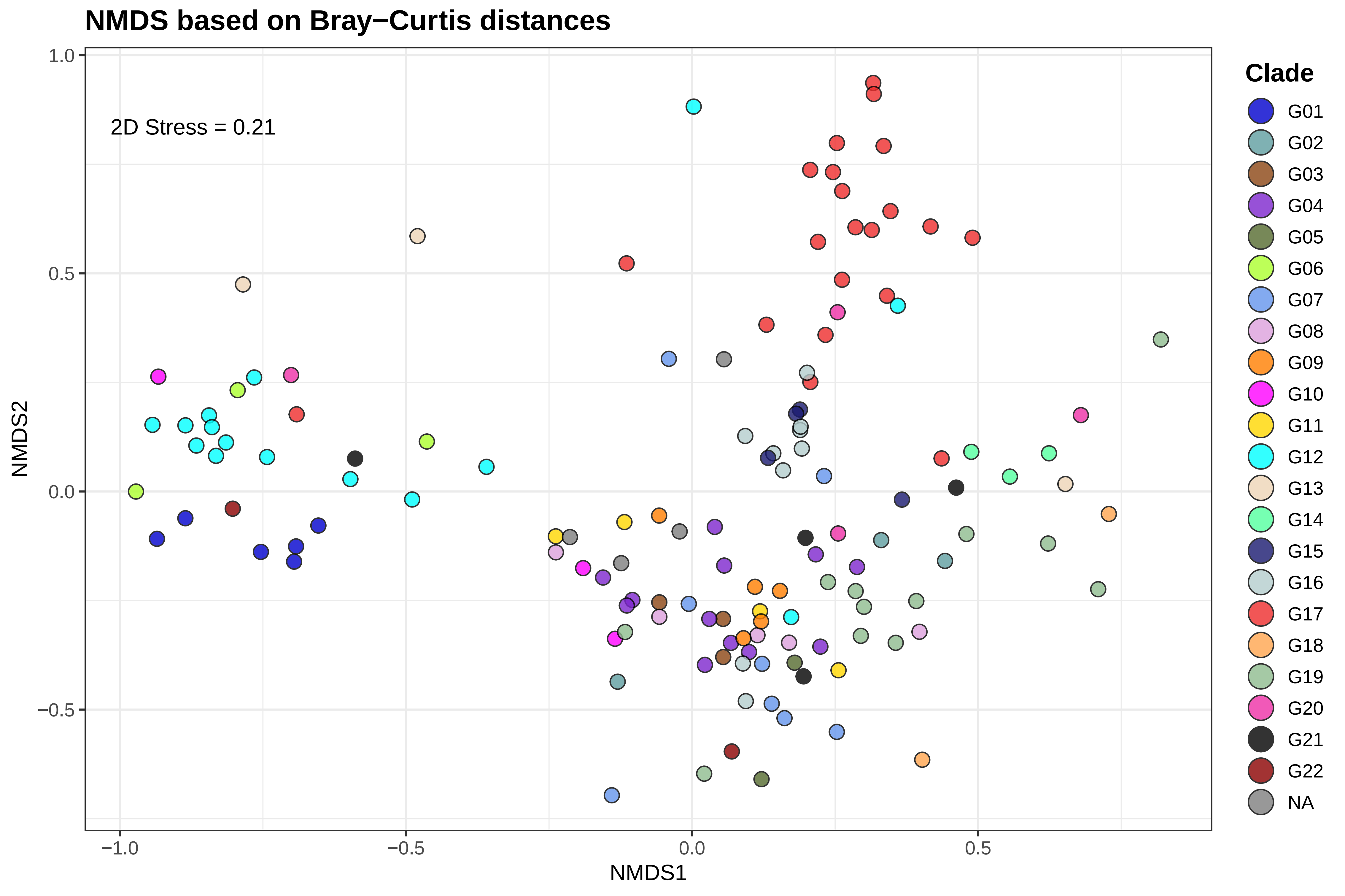

### Figure S3

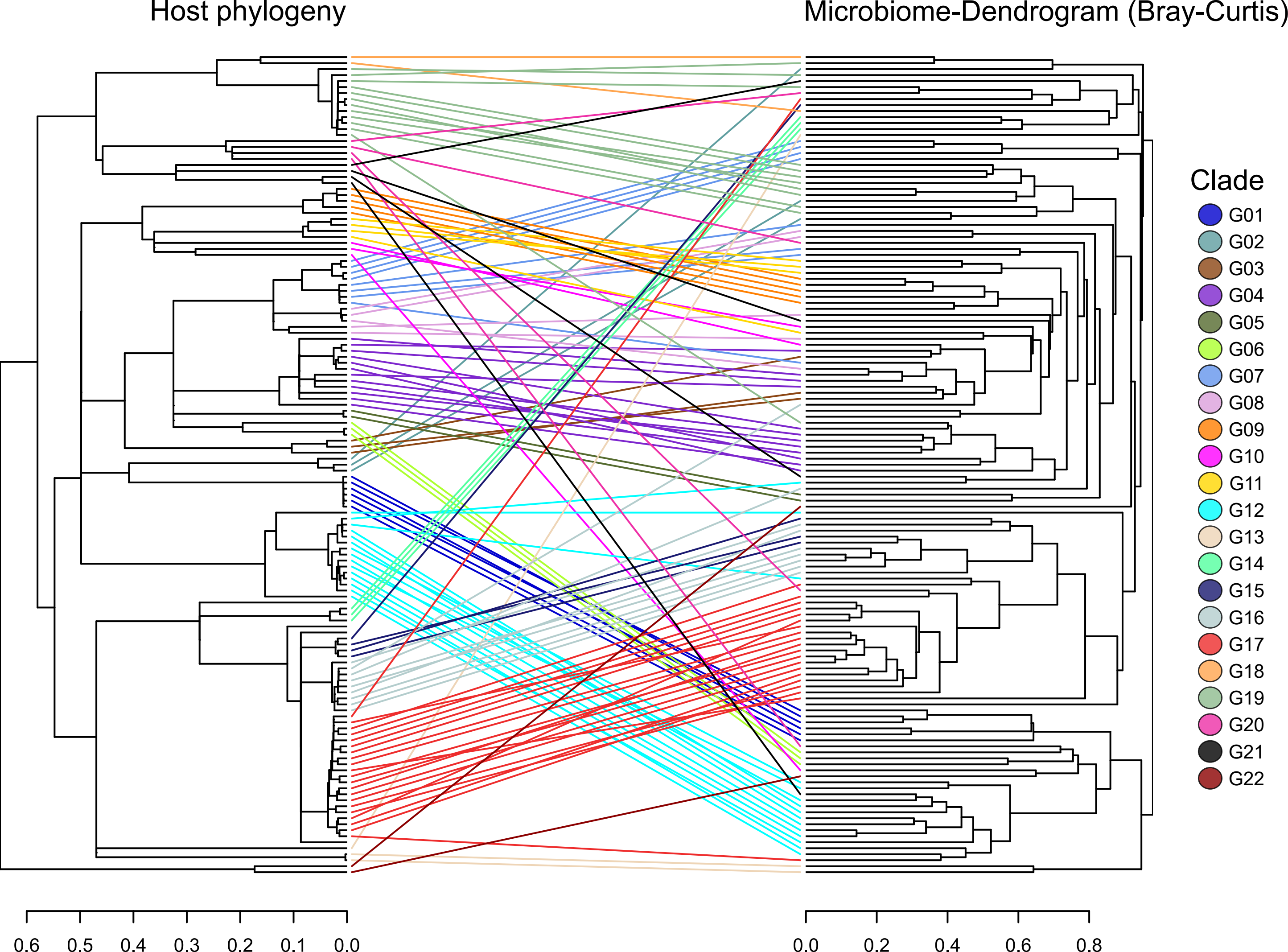
