## Supplementary Figure Legends for "Ecological Filtering is a Better Predictor of Microbiome Assembly than Coevolution in Marine Sponges"

**Supplementary Information**

**Figure S1.** Maximum-likelihood (ML) phylogeny of 144 Red Sea Haplosclerida based on a 25% concatenated alignment matrix of 1,153 Clade-Specific Element (CSE) loci (276,047 bp). Bootstrap probability (BP) values are shown at the nodes, with asterisk (*) indicating BP = 100. Scale bar is in substitutions per site. The coloured boxes represent the predefined clades (G01-G22) used for downstream analyses.

**Figure S2.** A nonmetric multidimensional scaling (NMDS) plot based on the Bray–Curtis dissimilarity of microbial communities found in different predefined Red Sea haplosclerid clades (see Figure S1).

**Figure S3.** A tanglegram, or co-phylogeny plot, linking the Red Sea haplosclerid phylogeny (1,153 Clade-Specific Element loci; 276,047 bp) on the left to the microbial dendrogram on the right. The coloured lines represent the predefined clades (G01-G22).
